# Spatial and temporal changes in [^3^H]JNJ-64413739 binding to purinergic P2X7 receptor (P2X7R) after status epilepticus induced by intracerebral kainic acid in the rat

**DOI:** 10.64898/2026.05.12.724505

**Authors:** Kristin H. Magnusdottir, Burcu A. Pazarlar, Cansu B. Egilmez, Jens D. Mikkelsen

**Affiliations:** Neurobiology Research Unit, University Hospital Copenhagen, Rigshospitalet, Copenhagen, Denmark; Physiology Department, Faculty of Medicine, Izmir Katip Celebi University, Izmir, Turkey; Department of Neuroscience, University of Copenhagen, Copenhagen, Denmark; Neurobiology Research, Department of Molecular Medicine, University of Southern Denmark, Denmark

**Keywords:** [^3^H]JNJ-64413739, P2X7, Neuroinflammation, Epileptogenesis, Autoradiography

## Abstract

Purinergic 2X7 receptor (P2X7R) is considered to play a critical role in neurological diseases, including epilepsy, and has also been proposed as a potential marker for neuroinflammation. This study aimed to validate the binding properties of the novel P2X7R radiotracer, [^3^H]JNJ-64413739, in rat brain using *in vitro* autoradiography, and additionally to explore spatial and temporal changes in P2X7R binding levels in a rat model of temporal lobe epilepsy using intrahippocampal administration of kainic acid (KA). Saturation of [^3^H]JNJ-64413739 to brain sections yielded a K_D_ of approximately 3 nM, with full saturation around 10 nM. The radiotracer was displaced with a structurally different P2X7R ligand, JNJ-47965567, indicating high affinity and specificity to rat P2X7R. In post epileptic rats, region-specific [^3^H]JNJ-64413739 binding revealed a bilateral increase in the hippocampal formation and its subregions few days after status epilepticus, peaking at day 30, and remained stable at this high level until day 90. Similar temporal profiles were identified in subcortical regions such as the thalamus. Interestingly, no change in binding was observed in the temporal and piriform cortices until day 30 where a dramatic increase occurred. Also, in the corpus callosum, significant increase was detected 30 days after the seizure. These results show that P2X7R binding, likely reflecting inflammation, is increased at delayed time points and exhibit region-specific patterns that is different from acute effects. Our findings suggest that P2X7R may contribute to sustained neuroinflammation and may be involved in those changes leading to epileptogenesis and the development of chronic epilepsy.

**Highlights:** [^3^H]JNJ-64413739 binds specifically to the purinergic P2X7 receptor (P2X7R) and saturates in the rat brain.

P2X7R binding increases in a region– and time-dependent manner following status epilepticus.

P2X7R binding remains elevated during chronic epilepsy in all examined brain regions.

P2X7R is considered a link between early seizures and sustained neuroinflammation and epileptogenesis.

## INTRODUCTION

Epilepsy is a common neurological disorder, affecting at least 70 million people globally (Banerjee *et al*., 2009; Singh and Trevick, 2016; Thijs *et al*., 2019; Waris *et al*., 2024). The disease is characterized by abnormal neuronal excitation, which results in the occurrence of spontaneous recurrent seizures (SRS) (Banerjee *et al*., 2009; Waris *et al*., 2024). Despite the symptoms are to a major extent medically treatable, approximately one-third of patients develop drug-resistant epilepsy (Chen *et al*., 2018; Kwan and Brodie, 2000), and the mechanisms underlying this resistance are poorly understood (Tang *et al*., 2017).

Clinical and experimental evidence suggests that neuroinflammation could play a role in the pathophysiology of epilepsy. In rodent models, status epilepticus (SE) induced by kainic acid (KA) rapidly increases the mRNA expression of pro-inflammatory cytokines, including interleukin-1β (IL-1β), IL-6, and tumor necrosis factor-α (TNF-α) in multiple brain regions, with IL-1β remaining elevated into the chronic phase (De Simoni *et al*., 2000; Jarvela et al., 2011; Minami *et al*., 1991; Minami *et al*., 1990; Pohlentz et al., 2022). Notably, increased IL-1β expression within microglia-like cells have also been reported in cortical and hippocampal tissues from patients with drug-resistant epilepsy (Choi *et al*., 2009; Leal et al., 2017; Ravizza *et al*., 2008). While this suggest that seizures induce neuroinflammation, inflammatory molecules may also heighten neuronal hyperexcitability, and lower seizure threshold, contributing to epileptogenesis (Aronica *et al*., 2017; Ravizza *et al*., 2011). The precise relationship between the timing when neuroinflammation develops and seizure propagation occurs as well as the specific brain regions where the processes originates and spread are not defined. This has also limited our understanding of the role of inflammatory processes in the development of epilepsy.

One transmitter, adenosine triphosphate (ATP), and its purinergic 2X7 receptor (P2X7R) are considered to play an important role in epilepsy (Andrejew *et al*., 2020; Sperlagh and Illes, 2014). P2X7R antagonists, including A-438079, brilliant blue G (BBG), and JNJ-4796556 have shown therapeutic potential in preclinical models of epilepsy (Engel *et al*., 2012; Jimenez-Pacheco *et al*., 2016).

P2X7R expression is upregulated at a protein level in the hippocampus within the first few hours after SE in mice induced by intra-amygdala KA injections (Engel *et al*., 2012; Huang *et al*., 2017; Morgan *et al*., 2020). Using P2X7R-EGFP reporter mice, the increase was found to take place in microglia and oligodendrocytes 72 hours after SE (Morgan *et al*., 2020). In the perspective of epileptogenesis, the level and function of P2X7R under the progression of epilepsy could be important. Epileptogenesis is the process by which a previously normal brain undergoes functional and morphological changes such as hippocampal atrophy, gliosis, axonal sprouting, and synaptic reorganization leading to the development of epilepsy (Engel and Pitkanen, 2020; Pitkanen *et al*., 2015; Sano *et al*., 2021), and these changes are important for the prognosis and treatment resistance (Fang *et al*., 2011; Thom, 2014). The aim of the present study was to evaluate P2X7R binding. In case this can be transferred to an in vivo setting using PET, P2X7 might be a potential biomarker of neuroinflammation Monitorization of P2X7R levels during all stages of epileptogenesis and in particular the transition from the latent period to epileptic state can provide valuable information into disease progression. However, spatial distribution of P2X7R under the development of epileptogenesis is lacking. Immunoblotting or gene expression is limited for comparing multiple brain regions and time points, whereas radiotracers and autoradiography enable both quantitative and qualitative estimation of receptor binding across many regions simultaneously.

Radioactive labelled JNJ-64413739 has been proposed as a radioligand with P2X7R selectivity (Kolb *et al*., 2019; Koole *et al*., 2019; Mertens *et al*., 2021; Mikkelsen *et al*., 2023). Due to significant species differences in P2X7R binding and pharmacological properties (Bartlett *et al*., 2014; Caseley *et al*., 2015), we first validated JNJ-64413739 as a radiotracer in the rat brain and found the tracer to be suitable at low nM concentrations with excellent specificity. We next investigated changes in the binding to P2X7R in various brain regions and at different time points under epileptogenesis and we here report major changes in P2X7R expression in a well-established model of chronic epilepsy (Levesque and Avoli, 2013; Rusina *et al*., 2021) during the acute, latent, and chronic phases of epileptogenesis and that these changes occur at different locations and at different times suggest P2X7R and the cells that express this receptor to be important in the interplay between neuroinflammation and epileptogenesis.

## EXPERIMENTAL METHODS

### Radioligand [^3^H]JNJ-64413739 and other materials

[^3^H]JNJ-64413739 ((*S*)-[^3^H](3-fluoro-2-(trifluoromethyl)pyridin-4-yl)(6-methyl-1-(pyrimi-din-2-yl)-1,4,6,7-tetrahydro-*5H*-[1,2,3]triazolo[4,5-*c*]pyridin-5-yl)-methanone) was synthesized and obtained from Tritec AG, Teufen, Switzerland (Mikkelsen *et al*., 2023). Specific activity of the radioligand was 81.4 Ci/mmol (3012 GBq/mmol). The radiochemical concentration was 1 mCi/mL on the day of synthesis. The specific activity left on the day of the experiment was determined for all experiments. The cold ligand itself JNJ-64413739, and JNJ-47965567 N-((4-(4-phenylpiperazin-1-yl)tetrahydro-2H-pyran-4-yl)methyl)-2-(phenylthio)nicotinamide were obtained as earlier described (Hopper *et al*., 2021; Mikkelsen *et al*., 2023).

### Animals, treatments, and handling of tissues

Adult male Sprague-Dawley rats (200-220 g BW) were obtained the animal facility of Ege University, Izmir, Turkey. All animals were housed in a controlled environment at 22 ± 2°C following a 12-hour light/dark cycle and had free access to standard food and water ad libitum. All animal procedures were approved by the Institutional Animal Care and Use Committee and were conducted in accordance with the United States Public Health Service’s Policy on Humane Care and Use of Laboratory Animals (Approval No. EGEHAYMER-2022–066).

Animals underwent anesthesia induced with ketamine (75–100 mg/kg i.p.) and xylazine (10 mg/kg i.p.) and placed in a stereotaxic frame. Reference points at the Bregma and Lambda were established, and the following coordinates were used for the injection into the right dorsal anterior hippocampus by using a rat brain atlas (Paxinos and Watson, 2006): Anteroposterior –2.04 mm; Mediolateral 1.10 mm; and Dorsoventral (DV) 3.81 mm. Animals were injected with 0.8 µg/2µl of kainate acid (1.9 mM) (Tocris, cat# 0222), dissolved in sterile saline. The injection was administered using a stereotaxic robot (Neurostar GmbH, Germany) with a 1.0-µl Hamilton micro-syringe equipped with a 30 G cannula at a flow rate of 1µl/min. To prevent solution backflow, the syringe remained in place for 5 minutes after the injection. Pre– and post– surgical care of the animals was conducted in accordance with the guidelines (Fornari *et al*., 2012). Following this procedure, the animals were placed in novel cages, where they were housed individually and monitored. Animals were killed by cervical dislocation under anesthesia, after which their brains were rapidly removed from the skull. The brains were gently snap-frozen in 2-methylbutane immersed in liquid nitrogen and stored at − 80°C until sectioning.

To construct a time-course study, animals were randomly divided into seven groups according to brain sampling points: acute (1,3,5 post day status epilepticus (pdSE)), latent (10, 30 pdSE), and chronic (60, 90 pdSE) (Levesque and Avoli, 2013; Rusina *et al*., 2021) with each group of animals taken at a given time point contained four to six animals, for a total of 38 animals. Additionally, control groups from acute and latent phase underwent sham operations with 2 μl of 0.9% isotonic saline solution following the same stereotaxic procedure, including a total of 25 animals. The total number of animals used in this study was 63.

The convulsive behaviors of the treated rats were scored based on the scale of 5 according to the Racine’s Scale (Racine, 1972). Rats were video-monitored for 10 hours during the light cycle over a 90-day period. Only those rats exhibiting at least 4-degree seizures within the first 4 hours post-treatment were included in the study. In addition to showing at least 4th-degree seizure, the rats exhibiting spontaneous recurrent seizures (SRSs) after one month were used for 60 and 90 pdSE groups. Only rats meeting these criteria were included in the longitudinal autoradiography study.

### In vitro autoradiography procedure for [^3^H]JNJ-64413739

Rat brains were sectioned into 16 µm thick coronal cryostat sections and mounted on Superfrost slides (Thermo Scientific). The sections were selected along the rostro-caudal axis according to different distances to the bregma and the regions of interest (ROIs). The sections were collected at approximately –3.6 mm to bregma, with the range of roughly –3.0 mm to –4.5 mm. To minimize tissue variability, experiments were conducted on adjacent sections, with a minimum of three technical replicates. The sections underwent a pre-incubation step twice for 10 minutes at room temperature in a 50 mM Tris-HCl buffer (pH 7.4) containing 0.5% bovine serum albumin (BSA). Following pre-incubation, the slides were incubated in an incubation buffer (50 mM Tris-HCl buffer (pH 7.4), 0.5% BSA, 5 mM MgCl2, 2 mM EGTA, and 6 nM [^3^H]JNJ-64413739 on a rotator at room temperature for 120 minutes.

For the saturation experiment, duplicate glass slides were incubated with [^3^H]JNJ-64413739 radioligand at concentrations ranging from 1 to 100 nM. Adjacent duplicate sections were incubated with the same concentration of the radioligand and nonspecific binding was determined in the presence of either 10 μM JNJ-47965567 or 10 µM JNJ-64413739.

For the displacement study, 12 nM [^3^H]JNJ-64413739 was mixed with JNJ-47965567 at concentrations ranging from 0.01 nM to 100 µM.

For the experimental study, representative sections from all animals were incubated together with 6 nM [^3^H]JNJ-64413739 in incubation buffer as described above and processed simultaneously.

The slides were washed twice in an ice-cold pre-incubation buffer for 10 minutes and briefly rinsed in an ice-cold distilled water, air-dried, and placed overnight in a paraformaldehyde chamber at 4°C. After fixation, the glass slides were air-dried and kept in a silica gel desiccator for 45 min to remove any leftover moisture. Glass slides were subsequently exposed to FUJI imaging phosphor plates for three days at 4 °C, alongside tritium standard ARC (American Radiolabelled Chemicals, Inc, USA) and tritium microscale Batch 21 A (GE Healthcare, UK). After exposure the imaging plate was then scanned using Amersham™ Typhoon™ IP Biomolecular Imager (GE healthcare, Chicago, USA) at pixel size 25 μm. Resulting autoradiograms were analysed with Image J software (Version 2.9.0, NIH).

### Data analysis and statistics

The ROIs were manually delineated for each tissue section using ImageJ based on landmarks in the individual section. Comparison to the rat brain atlas (Paxinos and Watson, 2006) were therefore considered tentative, as tissue sections from different animals may not in all cases completely overlap. Although this approach appears subjective, all measurements were independently performed by at least two independent researchers. Data analysis was performed in a blinded manner, ensuring that the nature of the animals was unknown for the researcher. Quantitative analysis of receptor binding was performed by measuring the mean optical density (OD) within each ROI. Standards with known concentrations were used to create a calibration curve, allowing interpolation of OD values from the ROIs to obtain corresponding radioactivity values (nCi/mg tissue equivalent (TE)). This interpolation was performed using the Rodbard curve function in ImageJ. The decay-corrected specific activity of the radioligand was used to convert nCi/mg TE to binding values (fmol/mg TE). Statistical analysis was performed using GraphPad Prism (version 10.4.0; GraphPad Software, San Diego, CA, United States). Specific binding was calculated by subtracting non-specific from the total binding.

One-way analysis of variance (ANOVA) followed by post-hoc Tukey test was performed for multiple comparisons of [^3^H]JNJ-64413739 binding levels between the sham treated control groups and KA-treated groups at various time points. A student t-test was performed to compare differences between ipsilateral and contralateral brain regions to the injection site. The error bars on the graphs represent the mean ± SD. Statistical significance was based on P-values below 0.05.

## RESULTS

### [^3^H]JNJ-64413739 binding to the rat brain: Saturation and displacement studies

Incubating rat brain sections with increasing concentrations of [^3^H]JNJ-64413739 and analysing binding intensity in the entire rat brain section revealed a concentration dependent saturation of specific binding. Co-incubation with either 10 µM of the cold tracer JNJ-64413739 (Fig 1A) or another structurally different small molecule previously shown to display affinity to P2X7, JNJ-47965567 (Fig 1B) showed that both molecules strongly inhibited the specific binding. Non-specific binding in the presence of both displacing substances revealed as expected a straight line in both cases (Figs 1A and B). The saturation experiment demonstrated full saturation at radiotracer concentrations around 10 nM, and incubation with higher concentrations of the tracer produced no more specific binding, whereas the total binding increase reflecting non-specific binding (Figs 1A, B).

**Figure 1:**
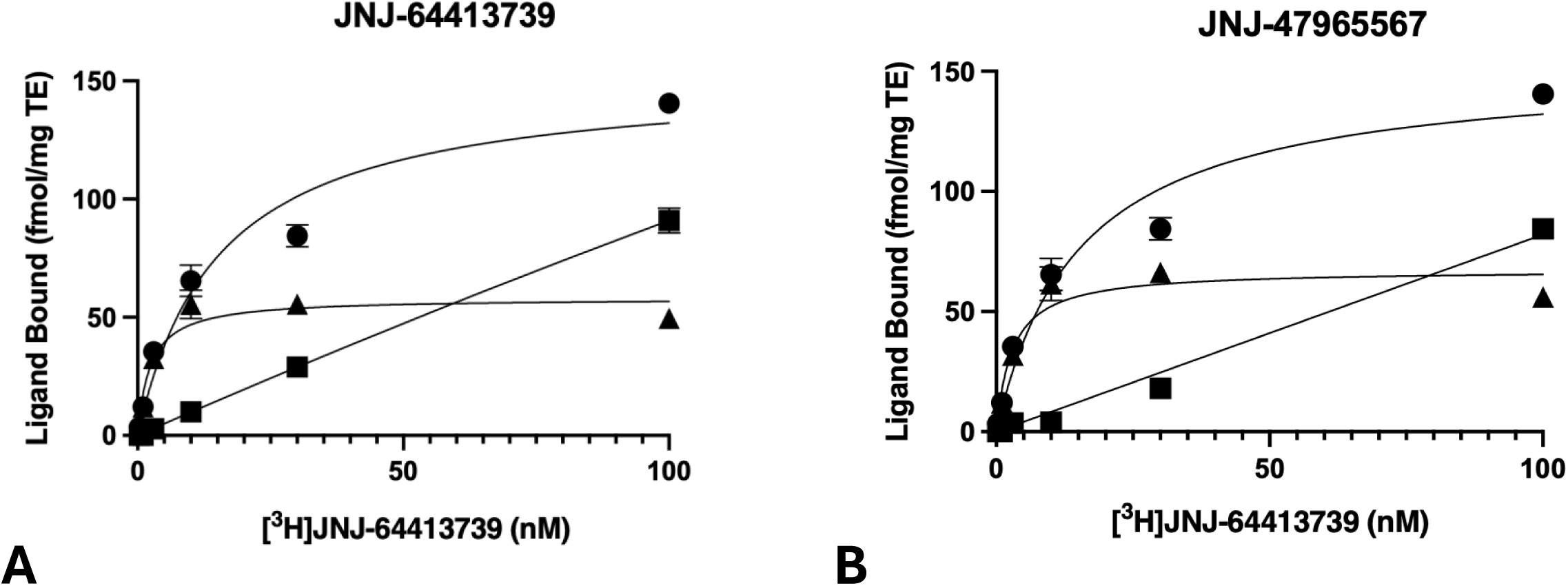
Saturation binding of [^3^H]JNJ-64413739 in the rat brain. Sections were incubated with increasing concentrations of the radioligand [^3^H]JNJ-64413739 with 10µM of either A) JNJ-64413739 or B) JNJ-47965567. In both cases, a saturation curve was demonstrated for the specific binding (▴), calculated to be the total binding (●) subtracted by non-specific binding (▪). The data represents the mean ± SD from two replicates per concentration point from one representative experiment where all sections were processed simultaneously.

Calculating the K_D_ from the saturation curves revealed a similar kinetics irrespective of the displacing ligand: K_D_ was determined to be 2.4 nM (95% CI: 1.4 nM-4.0 nM) after displacing with the cold ligand, and 2.9 nM (95% CI: 1.7 nM-5.0 nM) after displacement with JNJ-47965567.

Further, a displacement study was performed on other rat brain sections to assess binding specificity using increasing concentrations of the P2X7 antagonist, JNJ-47965567, along with 12 nM of [^3^H]JNJ-64413739 (Figs 2A and 2B). The ligand was able to reach full displacement at concentrations up to 1 µM. The calculated best fit IC_50_ was found to be 59.4 nM (95% CI: 44.9-79.2 nM) for JNJ-47965567 (Fig 2B).

**Figure 2:**
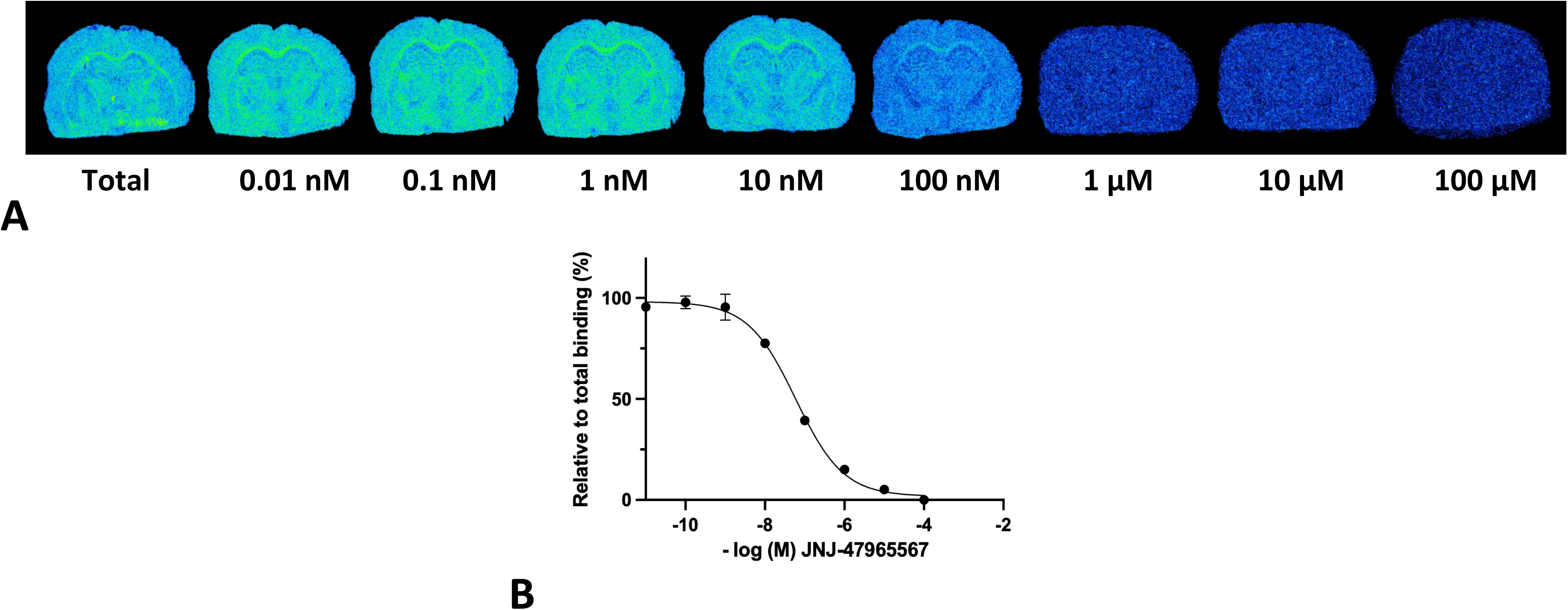
Representative autoradiograms from normal rat brain sections demonstrating displacement of [^3^H]JNJ-64413739 with increasing concentrations of JNJ-47965567. The labelling is relatively low and widely distributed throughout the brain also in white matter. B) The determination of binding levels for the entire section showed a complete block and calculated best fit IC_50_ was 59.4 nM (95% CI: 44.9-79.2 nM) for JNJ-47965567. Data represents the means from two replicates per data point.

### [^3^H]JNJ-64413739 binding in the intrahippocampal KA model

The binding levels of [^3^H]JNJ-64413739 to brain sections collected from rats exposed to intrahippocampal KA were examined across different time points after status epilepticus (post-day status epilepticus, pdSE). These time points spanned the acute (1, 3, 5 pdSE), latent (10, 30 pdSE) and into chronic phases (60, 90 pdSE) according to earlier definitions (Levesque and Avoli, 2013; Rusina *et al*., 2021). As illustrated in Figure 3, the autoradiographic images revealed major qualitative changes in density over the different time stages of epileptogenesis. The binding of [^3^H]JNJ-64413739 across the brain section exhibits a dynamic pattern with major changes dependent on time point and location. Across time points, the highest binding intensity observed one month post-injection during the latent phase in general. The binding was most prominent in the hippocampus and piriform cortex during this period, while in the chronic phase, the highest binding levels were seen in the corpus callosum (Figure 7). Control or sham groups (1, 3, 5, 30 days) were given intrahippocampal injections of saline. As there were no differences between control groups, the mean binding levels were pooled and used as the control baseline for all time points.

**Figure 3:**
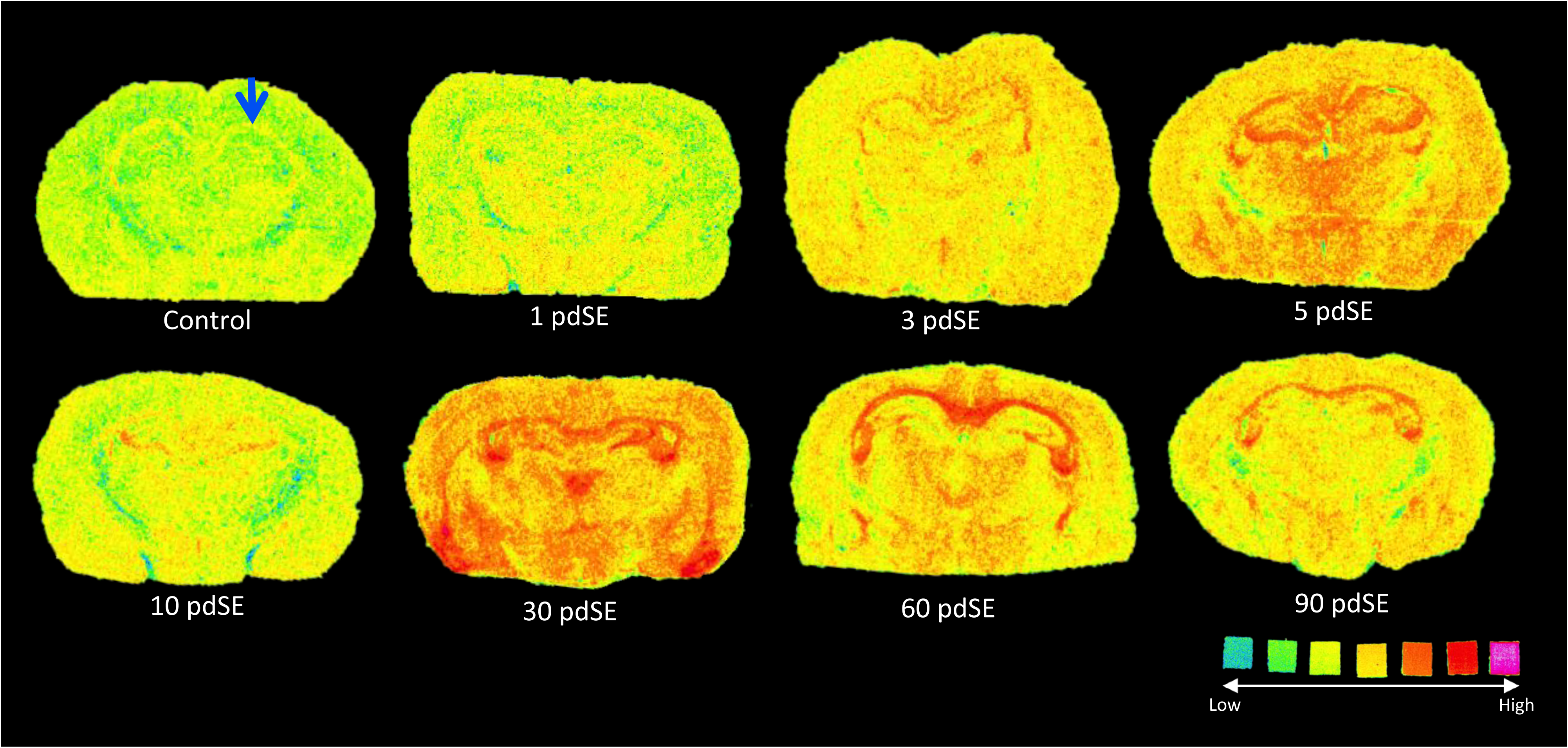
Representative autoradiograms of coronal sections from rat brains at the level of the injection site from rat brains injected intrahippocampally with KA into the right dorsal hippocampus (arrow). The individual brain sections are from rats taken at different time points after status epilepticus (post days after status epilepticus; pdSE). During the first days, the color intensity increases throughout the entire section, then decreases to baseline by day 10, and peaks on day 30. Note the very high binding intensity particularly in the temporal lobe, and in the corpus callosum.

### [^3^H]JNJ-64413739 binding in the hippocampus

The binding levels of [^3^H]JNJ-64413739 were analysed in the hippocampal formation and its subregions (CA1, CA3, and dentate gyrus). The temporal profile of binding levels was similar across all subregions, showing a gradual increase during the acute phase of epileptogenesis and gradually declining at 10 pdSE (Figs 4A-4D). The lowest binding levels were observed at this time point at the early latent phase, because the level increased again across all regions on day 30. At this time point the binding levels exceeded control values by >140% in the hippocampal formation, >220% in CA1, >170% in CA3, and >60% in the dentate gyrus. Within the chronic phase, the binding levels stabilize and perhaps slightly declined in particular in the CA1 region (Fig 4B). Thus, in the CA1 and CA3 a highly significant increase (90%) was observed after pdSE 10 (Figs 4B and 4C). The binding in the dentate gyrus remained more stable throughout both the latent and chronic phases (Fig 4D).

**Figure 4:**
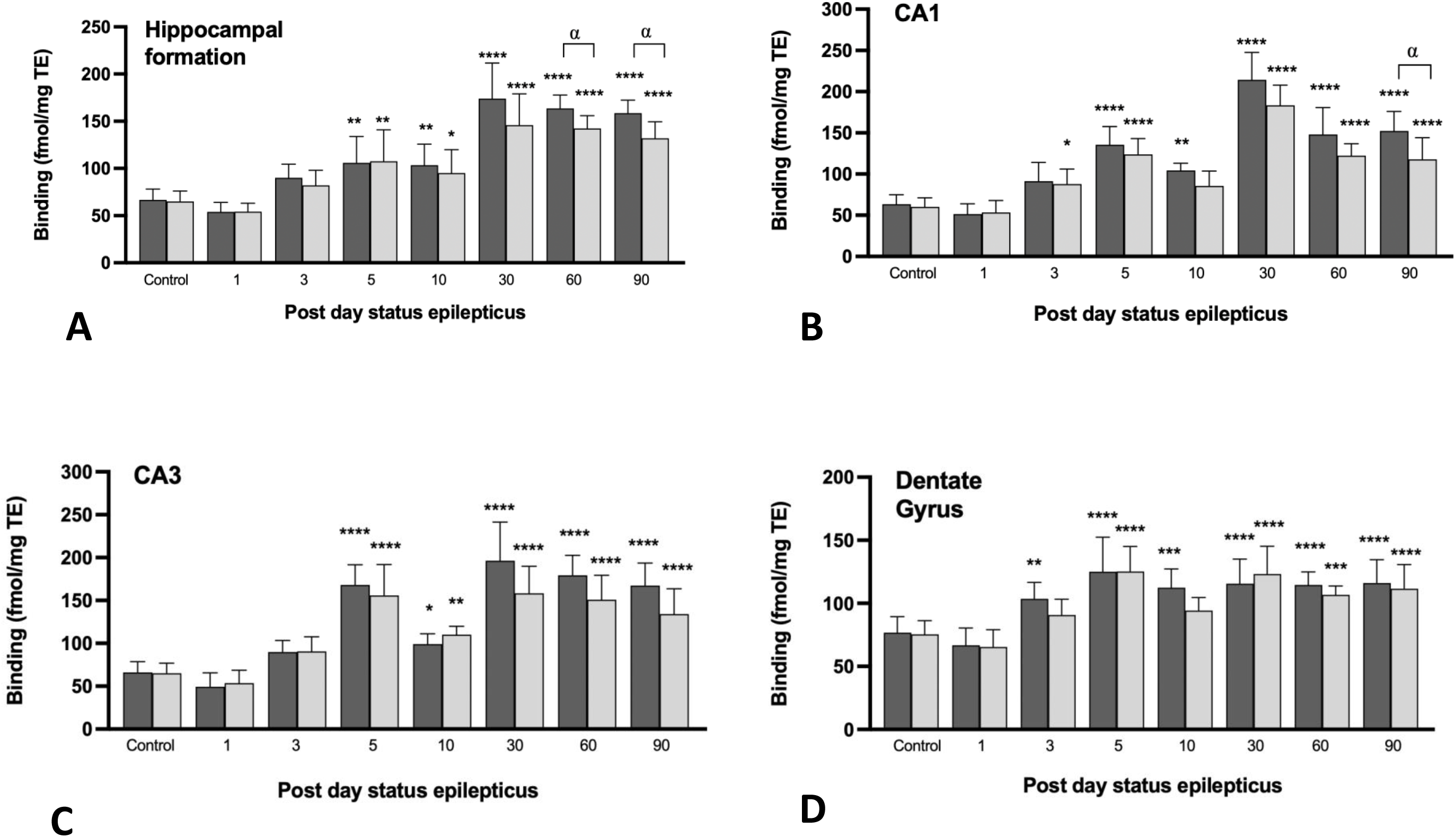
Quantitative analysis of [^3^H]JNJ-64413739 binding in the entire hippocampal formation (A), and in subregions CA1 (B), CA3 (C), and dentate gyrus (D). Measurements are performed on both ipsilateral (dark grey) and contralateral (light grey) sides. Statistical analysis was conducted using one-way ANOVA comparing treated animals with the control, demonstrating significant differences with *p < 0.05, **p < 0.01, ***p < 0.001, and ****p < 0.0001. For testing for lateralization, a Student’s t-test was conducted to compare the ipsilateral site and contralateral side, where “α” indicates statistical difference as seen only at late time points. The bars represent the mean ± SD.

Interestingly, as the only structure in the brain, a significant difference in binding was observed between the ipsilateral and contralateral sides of the injection site only during the chronic phase in the hippocampal formation (Fig 4A).

### [^3^H]JNJ-64413739 binding in the cerebral cortex

The binding levels of [^3^H]JNJ-64413739 in the temporal cortex increased slightly during the acute phase, followed by a decrease at 10 days post-SE reaching basal levels (Fig 5A). The binding intensity then increased significantly again one month after injection, with around a 2-fold increase compared to the control. This high binding intensity remains constantly high throughout the chronic phase (Fig 5A). The temporal profile of binding intensity in the cerebral cortex and to the hippocampal formation was similar. By contrast, the binding profile in the piriform cortex was much different to other cortical structures, as it remained relatively consistently low indistinguishable from basal levels until pdSE 30, where a highly significant and more than 2-fold bilateral increase compared to the control was observed in all animals (Fig 5B). Interestingly, after 30 pdSE the binding levels decreased dramatically back to basal levels in the ipsilateral hemisphere and remained at this intensity until 90 pdSE (Fig 5B).

**Figure 5:**
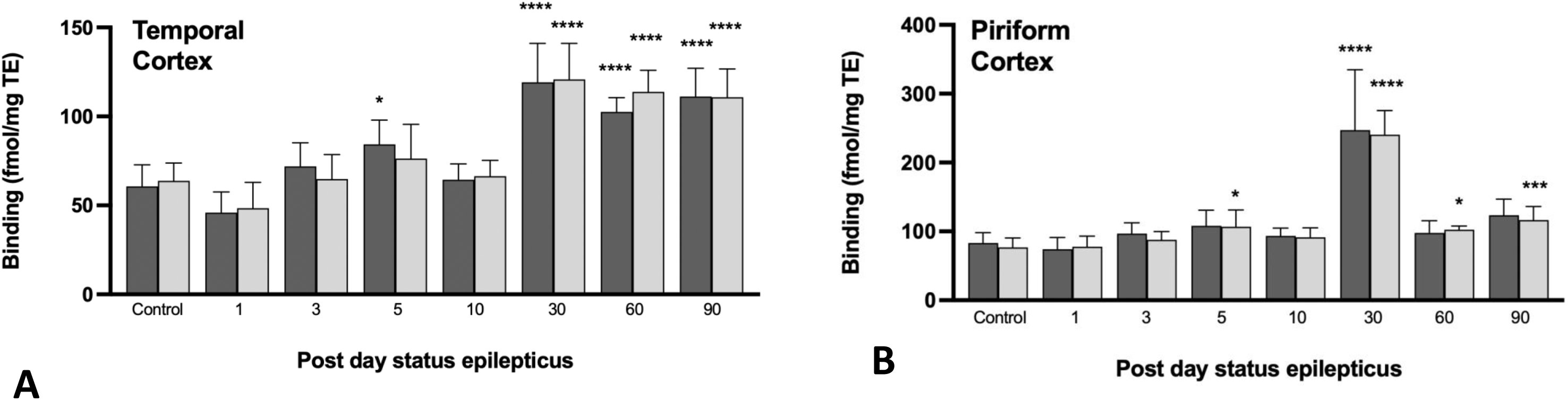
Quantitative analysis of [^3^H]JNJ-64413739 binding in the temporal cortex (A), and piriform cortex (B). Measurements are performed on both ipsilateral (dark grey) and contralateral (light grey) sides. Statistical analysis was conducted using one-way ANOVA comparing KA-treated animals with the control, demonstrating significant differences with *p < 0.05, **p < 0.01, ***p < 0.001, and ****p < 0.0001. The bars represent the mean ± SD.

### [^3^H]JNJ-64413739 binding in the subcortical regions

The binding levels of [^3^H]JNJ-64413739 in diencephalic structures such as the thalamus and hypothalamus increased throughout the acute phase, followed by a reduction on day 10 (Figs 6A and 6B). In the thalamus, the binding significantly increased by 80% on day 30 compared to the baseline control, and 110% compared to day 10 (Fig 6A). Binding intensity in the hypothalamus exhibited a similar pattern as the thalamus, though the magnitude was smaller (Fig 6B).

**Figure 6:**
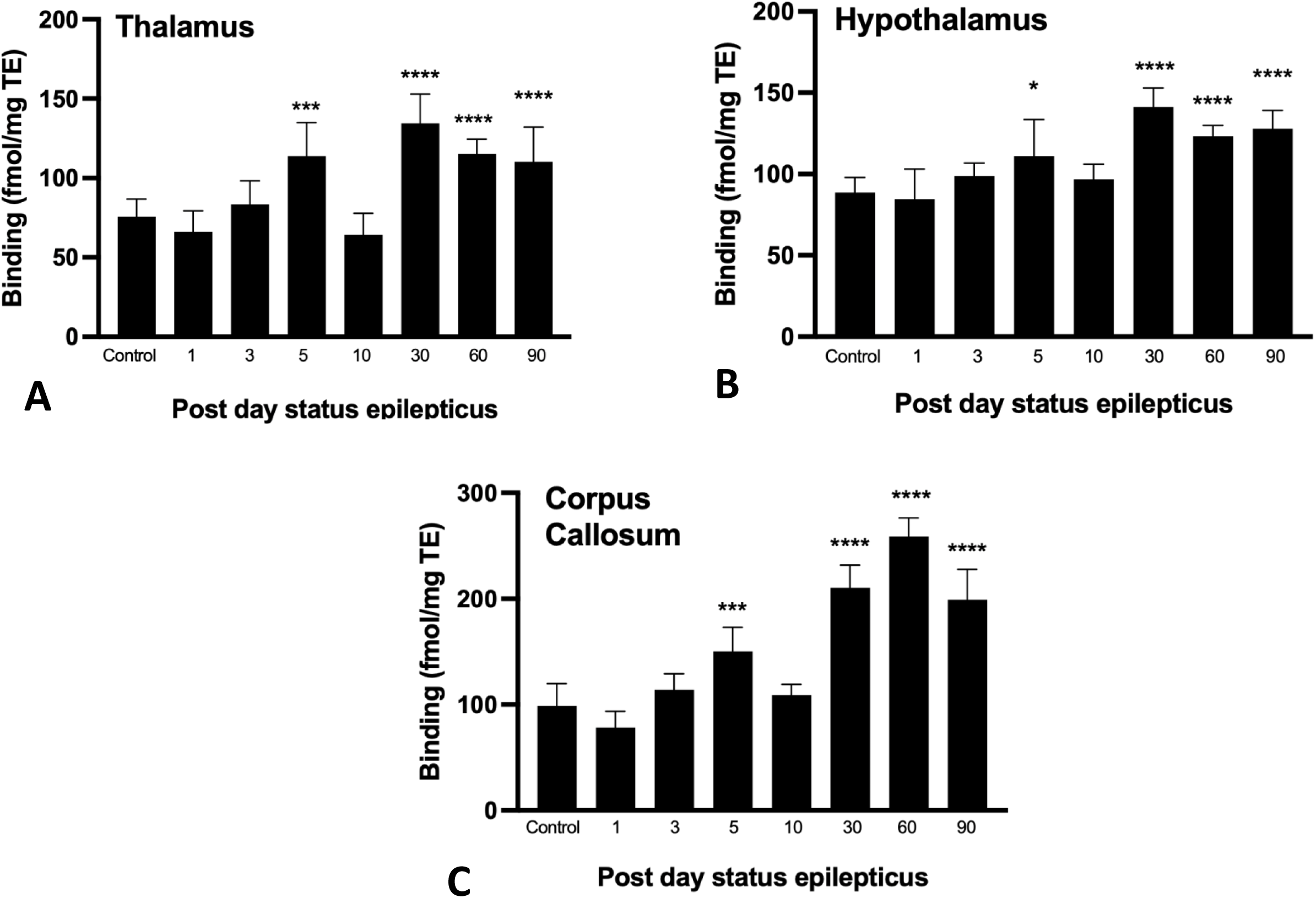
Quantitative analysis of [^3^H]JNJ-64413739 binding in the thalamus (A), hypothalamus (B), and corpus callosum (C) measured as the entire structures. Statistical analysis was conducted using one-way ANOVA comparing KA-treated animals with the control, demonstrating significant differences with *p < 0.05, **p < 0.01, ***p < 0.001, and ****p < 0.0001. The bars represent the mean ± SD

The corpus callosum showed a similar temporal pattern in binding over time, except that binding level continue to increase after pdSE 30 during the chronic phase, and first peaked 60 days after SE reaching a prominent increase of 160% compared to base line (Fig 6C).

## DISCUSSION

Given the significant species differences in P2X7 pharmacology (Bartlett *et al*., 2014; Caseley *et al*., 2015), we first validated the properties of [^3^H]JNJ-64413739 binding to normal rat brain sections using semi-quantitative autoradiography. The ligand was found to bind with high affinity and specificity to the P2X7R as confirmed by saturation and displacement experiments. The K_D_ was determined to be around 3 nM and full saturation was achieved at concentrations around 10 nM. Specificity was confirmed by full displacement of [^3^H]JNJ-64413739 by a structurally unrelated molecule, JNJ-47965567 at concentrations up to 1 µM. The specificity and affinity is validated and the binding results are similar to what we earlier obtained in human brain with the same radiotracer (Mikkelsen *et al*., 2023). Notably, it does not bind to mouse brain under the same conditions (unpublished observations) supporting interspecies differences. The same molecule has been radiolabeled with fluoride-18 and proposed as a biomarker for neuroinflammation in *in vivo* PET studies in the primate and human brain (Kolb *et al*., 2019; Koole *et al*., 2019; Mertens *et al*., 2021). Translating our findings into in vivo and PET, pharmacokinetics and blood-brain barrier penetration should be considered.

After confirming the tracer’s suitability, we next investigated changes in P2X7R binding across various brain regions and at different time points after KA-induced SE spanning from acute effects until chronic effects several months after. Unlike studies in human epileptic brain tissue, which are typically limited to the chronic phase of the disease, animal models offer us the opportunity to examine the full temporal change of P2X7R expression under epileptogenesis, including the early and latent phases. In contrast to gene expression analysis and immunoblotting, autoradiography allows for simultaneous screening of many regions and time points in the same animals in a single experiment. As illustrated in Figure 7, we present a detailed spatial distribution of P2X7R as revealed with radioligand binding to the receptor, and show major changes over time in different brain regions.

**Figure 7:**
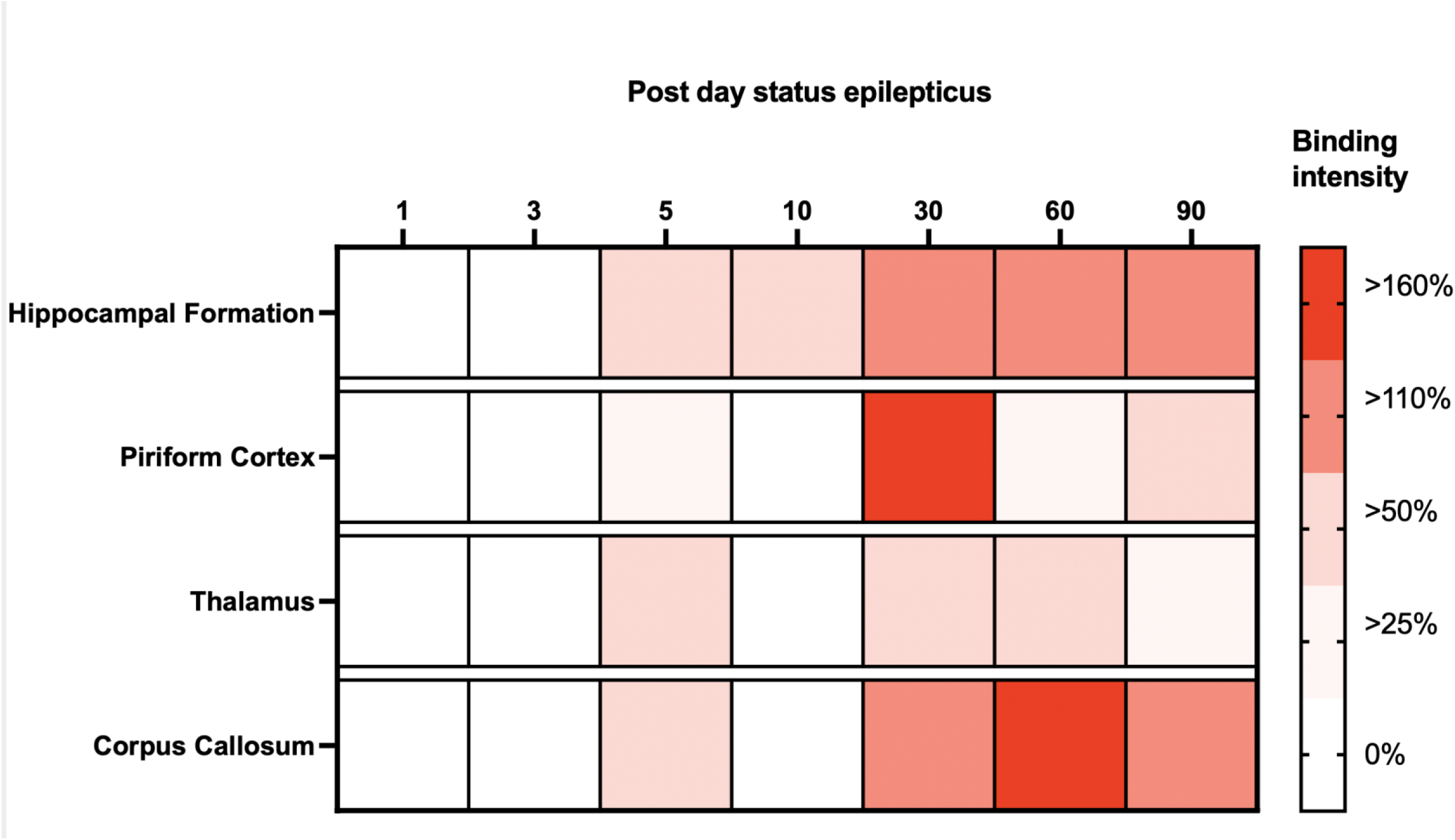
Heatmap representation of temporal changes in P2X7R binding across 4 selected brain regions: the hippocampal formation, piriform cortex, thalamus, and corpus callosum at various time points following SE. The intensity of red coloration reflects the percentage increase in P2X7R binding relative to control. The relative increase across regions and time illustrated in this heat map clearly show that changes in P2X7R expression occur differently in brain regions over time after SE.

Three main conclusions can be drawn from this work. First, P2X7R binding increased, not only at the site of the kainate injection, but throughout the brain within the first days after the seizures. This observation is in line with studies using different techniques to measure P2X7R expression. Earlier studies using immunodetection have also demonstrated an increase in P2X7R-immunoreactivity in the hippocampus within 48 hours after various types of seizures (Dona *et al*., 2009; Huang *et al*., 2017; Jimenez-Pacheco *et al*., 2013), but these data should be taken with some caution due to specificity of the antisera used. More recently, using a P2X7R-GFP reporter mice model, a significant increase in GFP-positive cells was observed in the CA1 and CA3 region of the hippocampus at 24– and 72-hours following SE (Morgan *et al*., 2020). At 72 hours, a twofold increase in GFP positive cells was seen, consistent with our autoradiographic findings demonstrating an increase in P2X7R binding in these same areas at the same magnitude 5 days after SE. Morgan et al. (2022) demonstrated that the increased expression took place in microglia and oligodendrocytes and not in neurons. Even radioligand binding was not detected at the cellular resolution, the dominant localization to the white matter is in accordance with a non-neuronal distribution.

Secondly, we observed that the hippocampal increase in P2X7R after this acute phase remained even up to 90 days after SE, which was the latest time point examined in the present study. The binding levels even increased in cortical and subcortical areas after the first days post-injection, and showed all a decline at 10 days after SE, and significantly elevated after this time point.

Thirdly, major changes in P2X7R binding were observed in the piriform cortex at a time point where spontaneous seizures begin to occur. P2X7R is known to play a crucial role in microglia activation and proliferation (Monif *et al*., 2009; Morgan *et al*., 2020), in both grey and white matter (Safaiyan *et al*., 2021; Woollacott *et al*., 2020). Furthermore, a rise in P2X7R binding was seen in the corpus callosum after 60 days, again suggesting changes that occur long time after the initial insult. The corpus callosum is a white matter region populated predominantly by oligodendrocytes but microglia and astrocytes are also present (Valerio-Gomes *et al*., 2018), supporting previous evidence of P2X7R expression in oligodendrocytes (Kaczmarek-Hajek *et al*., 2018) and suggesting that they may be involved in long-term effects of SE. Notably, these changes in white matter appeared to be specific for the corpus callosum, as other major myelinated trajectories such as the internal capsule was not affected.

Experimental evidence in rodents have shown that pro-inflammatory cytokines such as IL-1β, IL-6, and TNF-α mRNA expression increases in the cerebral cortex, hippocampus, thalamus, and hypothalamus within hours after SE (Eriksson *et al*., 2000; Jarvela et al., 2011; Lehtimaki *et al*., 2003; Minami *et al*., 1991; Minami *et al*., 1990; Plata-Salaman *et al*., 2000; Pohlentz et al., 2022). These are the same limbic brain regions in which we observed increased P2X7R binding during the first days after SE. Although P2X7R binding does not directly reflect cytokine expression or release, the overlap in spatial and temporal profile may suggest that, under the acute phase, increase in P2X7R binding occurs alongside increased TNFα and IL-1β expression (Vezzani *et al*., 1999). Also, increased IL-1β immunoreactivity has been observed in activated microglia in the hippocampus and parietal cortex following administration of KA, bicuculline, or pilocarpine during both acute and chronic phases (Ravizza *et al*., 2008; Vezzani *et al*., 1999). Notably, significant microglia activation and increased IL-1β expression within microglia-like cells have been reported in cortical and hippocampal tissues from patients with drug-resistant epilepsy (Choi *et al*., 2009; Leal et al., 2017; Ravizza *et al*., 2008). The presence of IL-1β expression may align with elevated P2X7R binding during the latent and chronic phases of epileptogenesis in the same regions, although their relationship was not examined and remains to be confirmed.

While the extracellular ATP concentration is typically under the threshold required for P2X7R activation under physiological conditions (Gil *et al*., 2022), it increases when cells are damaged to a concentration that likely activates the receptor. ATP binding through P2X7R promotes the activation and proliferation of microglia (Kaczmarek-Hajek *et al*., 2018), and regulates the nucleotide-binding domain, leucine-rich-containing family, pyrin domain-containing-3 inflammasome (Liao *et al*., 2025; Xie *et al*., 2014; Zheng *et al*., 2020). Priming via cytokines or lipopolysaccharide stimulation proceeding ATP activation of the P2X7R result in inflammasome assembly, caspase-1 activation, and the release of activated forms of inflammatory cytokines (Liu *et al*., 2017; Xie *et al*., 2014). This may increase neuronal hyperexcitability (Smith *et al*., 2023; Viviani *et al*., 2003), or even cause excitotoxicity (Di Virgilio *et al*., 1999; Hu *et al*., 2000; Viviani *et al*., 2003). Despite extracellular ATP concentrations under basal conditions are not enough to elicit effects via P2X7R (Beamer *et al*., 2021), our results now indicate that the number of receptor sites increase more than 2-fold in a given region and perhaps even more in individual cells, suggesting that ATP-P2X7R activation could be more important in chronic epilepsy than earlier believed.

P2X7R binding showed a spatial and temporal pattern that partially overlaps with Translocator Protein (TSPO), a well-established PET target for inflammation (Bouilleret and Dedeurwaerdere, 2021). In a rat model where pilocarpine was administered i.p. to induce SE, TSPO binding peaked between days 5 and 10 post-SE in the hippocampus, thalamus, piriform cortex, and amygdala, followed by a gradual decline over the next three months, eventually returning to baseline levels (Brackhan *et al*., 2016). Interestingly, in the piriform cortex, the peak occurred slightly later, around day 15-post SE (Brackhan *et al*., 2016). In our study, we observed a comparable initial increase in P2X7R binding during the first 10 days after SE. However, unlike TSPO, P2X7R binding returned to baseline levels on day 10 in most regions, except the hippocampus. Notably, P2X7R binding reached a peak around day 30 across all brain areas examined, except the corpus callosum, where the peak occurred at day 60. These findings suggest that the inflammatory response involving P2X7R is prolonged compared to TSPO. Comparison to TSPO binding is interesting, because TSPO is the most common neuroinflammation target (Bouilleret and Dedeurwaerdere, 2021). There are misinterpretations and limitations in the use of TSPO (Kreisl *et al*., 2020), and our data suggest that P2X7R may be detecting other changes under neuroinflammation and could be an alternative target for evaluating neuroinflammation.

In summary, this study provides a detailed time course of P2X7R binding throughout all phases of epileptogenesis in rats, and shows that P2X7R levels are sustained also in the chronic phase (see Fig 7). Furthermore, we detect P2X7R binding in the corpus callosum, a white matter region with a non-neuronal distribution. Finally, the very large increase in P2X7R binding at the interface between the latent and chronic phase indicates the P2X7R is a marker and perhaps a regulator of permanent changes in the network involved in chronic epilepsy and recurrent spontaneous seizures. Together, these findings show prolonged changes in P2X7R binding following epileptic insults but the consequences for neuroinflammation and eventual disease progression have not been ruled out.

## CONSENT FOR PUBLICATION

All authors have approved the final draft and publication of the present manuscript

## AVAILABILITY OF DATA AND MATERIALS

All data are available on request to JDM

## AUTHOR CONTRIBUTIONS

BAP and JDM designed the research and interpreted the initial results. BAP and CBE conducted the animal surgery and observations, and BAP conducted the binding experiments. BAP and KHM conducted the data analysis and interpreted the results. KHM and JDM wrote the draft of the manuscript, and all authors revised and approved the final draft of the manuscript.

## Abbreviations

ATP: Adenosine 5’-triphosphate
BSA: Bovine serum albumin
GABA: γ-aminobutyric acid
KA: Kainic acid
OD: Optical density
pdSE: Post day status epilepticus
P2X7R: Purinergic P2X7 receptor
ROI: Region of interest
SE: Status epilepticus
SRS: spontaneous recurrent seizures

